# Membrane tension spatially organizes lysosomal exocytosis

**DOI:** 10.1101/2022.04.22.489160

**Authors:** Hugo Lachuer, Laurent Le, Sandrine Lévêque-Fort, Bruno Goud, Kristine Schauer

**Affiliations:** Cell Biology and Cancer Unit, Institut Curie, PSL Research University, Sorbonne Université, CNRS UMR144, 75005 Paris, France; Université Paris-Saclay, CNRS, Institut des Sciences Moléculaires d’Orsay, 91405, Orsay, France; Tumor Cell Dynamics Unit, Inserm U1279, Gustave Roussy Institute, Université Paris-Saclay, 94800 Villejuif, France

## Abstract

Lysosomal exocytosis is involved in many key cellular processes but its spatio-temporal regulation is poorly known. Using total internal reflection fluorescence microscopy (TIRFM) and spatial statistics, we observed that lysosomal exocytosis is not random at the adhesive part of the plasma membrane of RPE1 cells but clustered at different scales. Although the rate of exocytosis is regulated by the actin cytoskeleton, neither interfering with actin or microtubule dynamics by drug treatments alters its spatial organization. Exocytosis events partially co-appear at focal adhesions (FAs) and their clustering is reduced upon removal of FAs. Changes in membrane tension following a hypo-osmotic shock or treatment with methyl-β-cyclodextrin was found to increase clustering. To investigate the link between FAs and membrane tension, cells were cultured on adhesive ring-shaped micropatterns, which allows to control the spatial organization of FAs. By using a combination of TIRFM and fluorescence lifetime imaging microscopy (FLIM) microscopy, we revealed the existence of a radial gradient in membrane tension. By changing the diameter of micropatterned substrates, we further showed that this gradient as well as the extent of exocytosis clustering can be controlled. Together, our data indicate that the spatial clustering of lysosomal exocytosis relies on membrane tension patterning controlled by the spatial organization of FAs.

## Introduction

Exocytosis is an evolutionary novelty shared among all eukaryotes (i.e. a synapomorphy) (Kloepper et al., 2007). It relies on the SNARE machinery probably inherited from an archaeal ancestor (Neveu et al., 2020). In addition to Golgi-derived vesicles along the secretory pathway, late endosomes, lysosomes and lysosome-related organelles also undergo exocytosis. Lysosomal exocytosis (referred to the secretion from late endosomes/lysosomes) is involved in the secretion of enzymes (Samie and Xu, 2014) and exosomes (Kowal et al., 2014). It supports plasma membrane repair (Andrews and Corrotte, 2018) as well as the remodeling of the microenvironment. Besides these general functions, lysosomal exocytosis fulfills specific roles in several cell types, such as the growth of neurites (Arantes and Andrews, 2006), axonal myelination (Chen et al., 2012), cell communication through ATP release in astrocytes (Dou et al., 2012), pseudopode formation in phagocytosis (Huynh et al., 2007), secretion of cytotoxic granules in lymphocytes (Peters et al., 1991), MHC-II antigen presentation (Geuze, 1998) and bone resorption in osteoclasts (Lacombe et al., 2013). It has a fundamental importance in several pathological contexts. For instance, lysosomal exocytosis is exploited by some β-coronavirus for their egress (Ghosh et al., 2020; Chen et al., 2021) or by cancer cells to enhance invasion through the secretion of metalloproteases, especially at invadopodia (Hoshino et al., 2013; Machado et al., 2015). Importantly, impairment of lysosomal exocytosis has been implicated in lysosomal storage disorders (LSDs) (LaPlante et al., 2006). Enhancing lysosomal exocytosis to release undigested lysosomal contents is a promising therapeutic strategy in these diseases (Samie and Xu, 2014).

Seminal work on the molecular machinery maintaining exocytosis, particularly vesicular (v-) and target (t-) SNAREs that are critical for the fusion of secretory vesicles arriving at the plasma membrane (PM) (Novick et al., 1980; Söllner et al., 1993; Fernández-Chacón et al., 2001), uncovered key mechanisms that regulate the frequency of secretory events (Gundelfinger et al., 2003; Kasai et al., 2012). Exocytosis has been known to be polarized toward active zones in neuronal cells for a long time (Südhof, 2012). In recent years, the question has arisen where exocytosis takes place and how cells regulate secretion at specific cellular sites in non-neuronal cells. After some conflicting results (Schmoranzer et al., 2000; Keller et al., 2001), it has been now clearly demonstrated that exocytosis is not random but clustered even in non-polarized cells (Sebastian et al., 2006; Yuan et al., 2015; Urbina et al., 2018; Fu et al., 2019). Evidence also exists that lysosomal intracellular positioning is non-random (Schauer et al., 2010; Ba et al., 2018). Moreover, in polarized epithelial cells, lysosomal exocytosis is targeted to the basolateral membrane (Xu et al, 2012). However, the spatial regulation of exocytosis in non-polarized cells and its mechanisms have not been explored.

Lysosomal exocytosis relies on VAMP7 (Martinez-Arca et al., 2000; Proux-Gillardeaux et al., 2007; Verderio et al., 2012), a v-SNARE that is insensitive to tetanus and botulinum neurotoxins (hence its other names TI-VAMP for Tetanus neurotoxin Insensitive Vesicle-Associated Membrane Protein) (Galli et al., 1998). In epithelial cells, VAMP7 interacts with the t-SNAREs syntaxin (STX) 3 (Vogel et al., 2015) and STX4 (Williams et al., 2014) found at the PM and with STX7. Whereas STX3/4 are involved in exocytosis, STX7 is only involved in intracellular endosomal fusion events (Ward et al., 2000; Wade et al., 2001; Bogdanovic et al., 2002).

In the present study, we use tools from spatial statistics to analyze live imaging data of fluorescent VAMP7 obtained by TIRFM on RPE1 cells undergoing lysosomal exocytosis. We report that spatial organization of lysosomal exocytosis is regulated by membrane tension gradient that relies on the spatial distribution of FAs.

## Results

### 1) Exocytosis from lysosomes is not random

To monitor lysosomal exocytosis, we transfected RPE1 cells with a pHluorin construct of the v-SNARE VAMP7, VAMP7-pHluorin. The pHluorin signal is quenched in the lumen of acidic vesicles but unquenched during exocytosis when protons from the lysosomal lumen are released (F1A). Using dynamic TIRFM imaging, we manually detected exocytosis events characterized by a sudden increase in intensity followed by a decay clearly visualized in kymographs (F1B). The decrease in intensity corresponds to the 2D diffusion of VAMP7-pHluorin in the plane of the PM. Its diffusion kinetic can be fitted by a single decreasing exponential function with a half-life of about 1.69 ± 0.83s (S1A-S1B). Detected exocytosis can be represented by an intensity map where the intensity λ represents the local expected number of event/μm^2^ (F1C). RPE1 cells display a high lysosomal exocytosis rate as several hundred exocytosis events in a typical cell could be observed within 5min, corresponding to an exocytosis rate of 39×10^−5^ ± 31×10^−5^ exocytosis/μm^2^/s (F1D). VAMP7 has been reported as a marker of the endosomal/lysosomal compartments. However, it substantially colocalizes with Golgi-derived vesicles in some cell types (Chaineau et al., 2009), especially in neuronal cells (Burgo et al., 2012). To confirm that VAMP7 specifically marks lysosomal exocytosis in RPE1 cells, we treated cells with Golgicide A, an inhibitor of the Arf1 GEF (Guanine nucleotide Exchange Factor) GBF1, which induces Golgi apparatus dispersion and inhibits the Golgi-derived vesicles secretion (Saenz et al., 2009). As shown in (F1E), Golgicide A did not significantly alter the exocytosis rate. Contrary, inhibition of the lysosomal V-type ATPase using Bafilomycin A1 significantly reduced the exocytosis rate (F1F). Lysosomal exocytosis can be stimulated by histamine through the G1q-PKC pathway (Verweij et al., 2018). As expected, treating RPE1 cells with histamine significantly increased the exocytosis rate (F1G) immediately after the addition of histamine (S1C). Taken together, these results demonstrate that VAMP7 exocytosis represents *bona-fide* lysosomal exocytosis in RPE1 cells.

The obtained exocytosis maps can be visualized as patterns of points (F1C). Such patterns can be the result of an uniformly random process (Complete Spatial Randomness, CSR) or reflects either clustering (i.e. aggregation) or dispersion (i.e. ordering with an inhibition surface around each point of exocytotic event) (Diggle, 1983; Lachuer et al., 2020). We obtained a large dataset of 183 cells showing 32 880 exocytosis events to test the CSR hypothesis and to characterize the spatio-temporal properties of lysosomal exocytosis. The observed spatio-temporal characteristics of exocytosis were compared to CSR Monte-Carlo simulations (see methods). First, we found that exocytosis events present a smaller nearest-neighbor distance (NND) than expected in the case of CSR, which indicates clustering at the scale of immediate neighbors (F1H). Moreover, events are more distant to cell borders than expected demonstrating that exocytosis is much less frequent close to cell borders (F1I). Events are also anisotropically distributed in the cells *i*.*e*. have a preferential direction (F1J). To explore this anisotropy, we seeded cells on rectangular micro-patterns forcing the orientation of cells into two possible directions (left or right) (S1D). Results confirmed anisotropy in exocytosis and showed that the secretory direction correlates with the Golgi-Nucleus axis (S1D).

The most pertinent tool to analyze a spatial structure is the Ripley’s K function that measures the average number of neighbor events at a given scale. The average curve obtained from observed events significantly deviated from the expected one in case of CSR, indicating clustering of lysosomal exocytosis, even at the scale of several micrometers (F1K). The CSR envelope is not symmetric around 0, reflecting a slight bias introduced by the boundary corrections in the computation of the Ripley’s K function. Note that the treatment with Golgicide A, Bafilomycin A1 and histamine did not change the spatial organization of exocytosis (S1E-G), indicating that the rate and spatial patterning of exocytosis are independent features. We noticed a slight correlation between exocytosis rate and clustering (S1H). This correlation is not due to a bias in the measure, because Ripley’s K function is independent on the number of events (Baddeley et al., 2015) and indeed, conserving an arbitrary percentage of the recorded events does not affect the Ripley’s K function (S1I). We conducted a similar analysis for the temporal distribution of exocytosis events. The temporal Ripley’s K function demonstrates a temporal clustering (S1J). However, a Fourier analysis revealed that this clustering is not due to a periodicity in the exocytosis rate (S1K). Lastly, we quantified the coupling between spatial and temporal dimensions using the spatio-temporal Ripley’s K function (S1L), which provides information about the independency of the temporal and spatial coordinates. This analysis revealed that a substantial proportion of cells presents a spatio-temporal coupling and among cells with a significant coupling, 82.7% have a positive coupling *i*.*e*. events that are close in space are also more likely to be close in time. Together, our analysis confirmed that lysosomal exocytosis is a non-random process in space and time.

### 2) Lysosomal exocytosis is coupled to internal focal adhesions in a cytoskeleton-independent manner

To investigate the mechanisms underlying the spatial clustering of lysosomal exocytosis events, we first tested whether the t-SNAREs interacting with VAMP7 show clustering, as proposed previously (Xu et al., 2012). Therefore, we analyzed the spatial patterns of STX3 and STX4 (S2A-C). Although STX3/4 presented a significant clustering, it was much weaker than the one of exocytosis events, and did not recapitulate several features of VAMP7 exocytosis such as low frequency close to cell borders and a short scale clustering. Next, we focused on FAs, shown to be targeted by a subpopulation of lysosomes that were characterized to be the MAPK scaffoled complex p14-MP1 (p14-MP1+) (Schiefermeier et al., 2014) as well as by Golgi-derived RAB6+ vesicles (Fourriere et al., 2019). We quantified the co-appearance of VAMP7-pHluorin with FAs using the FA protein paxillin (Paxillin-mCh) as a marker in co-transfected cells (F2A-B). The co-appearance index was significantly higher than expected from CSR Monte-Carlo simulations (F2B). Interestingly, VAMP7 lysosomal exocytosis only appeared at internal FAs, consistent with the observation that exocytosis frequency is low close to cell borders (F2A). To further test the role of FAs, we cultured cells on Poly-L-Lysine (PLL) substrates to inhibit the formation of FAs (S2D). Under this condition, the clustering of lysosomal exocytosis decreased significantly (F2C), indicating a role of FAs in the spatial organization of exocytosis. We noticed that exocytosis rate was significantly enhanced on PLL substrate, probably due to a smaller adhesive surface under this condition (S2E).

FAs are closely linked to both microtubules (MT) (+) ends (Stehbens and Wittmann, 2012; Stehbens et al., 2014; Seetharaman and Etienne-Manneville, 2019), and the cortical acto-myosin cytoskeleton (Miklavc and Frick, 2020). Surprisingly, MT depolymerization by nocodazole treatment did not reduce clustering (F2D) nor the exocytosis rate (F2E). Depolymerisation of F-actin by a low dose of cytochalasin D significantly reduced the exocytosis rate (F2G), however did not reduce lysosomal clustering (F2F). This suggested a role of actin in facilitating fusion of lysosomes with PM but not in the organization of exocytosis patterning. Moreover, myosin-2 (MYH9) inhibition by para-nitro-blebbistatin treatment affected neither exocytosis clustering nor exocytosis rate (F2H-I). Taken together, these results demonstrate that FAs regulate lysosomal exocytosis patterns in a cytoskeleton-independent manner. Moreover, they confirm a targeting of lysosomal secretion events to internal FAs and exclusion from external ones.

### 3) Exocytosis clustering depends on membrane tension

We tested the role of physical parameters, such as membrane tension, known to regulate the exocytosis rate (Gauthier et al., 2011; Wen et al., 2016; Kliesch et al., 2017; Shi et al., 2018; Wang and Galli, 2018; Wang et al., 2018). We applied a hypo-osmotic shock and monitored VAMP7 exocytosis 15min after the shock. Hypo-osmotic shock causes cell swelling leading to an increased membrane tension. VAMP7+ exocytosis has been reported to be less frequent after hyper-osmotic shock (Wang et al., 2018) and hypo-osmotic shock decreased exocytosis rate in experimental and theoretical models (Zwiewka et al., 2015; Mao et al., 2021). Surprisingly, the hypo-osmotic shock significantly reduced the exocytosis rate in RPE1 cells (F3B). However, hypo-osmotic shock significantly increased the clustering of lysosomal exocytosis (F3A). It also increased the co-appearance between exocytosis events and FAs (F3C). To further confirm these results, we treated cells with methyl-β-cyclodextrin that depletes cholesterol from the PM, and thus has been proposed to affect membrane tension (Hissa et al., 2017; Biswas et al., 2019; Cox et al., 2021). Cyclodextrin treatment significantly increases clustering, similarly to hypo-osmotic shock treatment (F3D). Moreover, cyclodextrin treatment also decreases exocytosis rate (F3E) and increases the co-appearance between exocytosis events and FAs (F3F).

Next, we directly measured membrane tension using the Fluorescence Lifetime Imaging Microscopy (FLIM) probe Flipper-TR (Colom et al., 2018). The Flipper-TR intercalates in membranes, where the membrane microenvironment favors one of two possible molecular conformations of the Flipper-TR (planar or orthogonal). The two conformations display different fluorescence lifetimes. Higher local membrane tension leads to higher fluorescence lifetimes. We quantified fluorescence lifetime in living cells using a TIRF-FLIM device under different experimental conditions. We observed that the membrane tension at the ventral part of the cell is not homogeneous in cells seeded on fibronectin-coated surface. Interestingly, we found that membrane tension presents variability compatible with the micrometer scale of exocytosis clustering (F3G). Surprisingly, no major variation in the Flipper-TR fluorescence lifetime was observed following a hypo-osmotic shock or treatment with cyclodextrin (S3A-B). Because lysosomal clustering is regulated by FAs, we next quantified fluorescence lifetime in living cells seeded on PLL-coated surface. Despite their difference in exocytosis clustering, the average lifetime was again similar in these two conditions (F3H). Together, these results indicate that global membrane tension does not regulate clustering of exocytosis. However, cells grown on fibronectin showed clustering of similar lifetime values at several regions, whereas these values appeared more homogeneously distributed in cells on PLL (F3G). Thus, we measured the spatial auto-correlation using the Moran’s index, which increases in case of clustering of pixels values. Moran’s index was indeed significantly higher in cells plated on fibronectin than on PLL (F3I). This suggests that the spatial organization of lysosomal exocytosis is controlled by regional heterogeneity in membrane tension rather than the global tension at the whole cell level. Moreover, our results indicate that the presence of FAs favors a compartmentalization of membrane tension.

### 4) Intracellular coupling between exocytosis probability and membrane tension

A simultaneous observation of the Flipper-TR signal and exocytosis events is technically very challenging. Therefore, to correlate membrane tension and exocytosis events, we normalized RPE1 cell geometries using adhesive ring-shaped micropatterns. An advantage of micropatterned cells is the possibility to standardize the adhesive surface and FA distribution. On ring-shaped micropatterns (F4A), FAs are formed at the inner and outer borders of the ring mimicking the inner and peripheral FAs found in non-patterned cells (S4A-B). Using the Ripley’s K function, we found that the clustering of lysosomal exocytosis also occurs in patterned cells, although significantly weaker than in non-patterned cells (F4B). Plotting the radial average density of exocytosis events demonstrated that cells exhibit an enrichment of events at half of the cell radius as the density deviated there from the expected CSR case (F4C). Moreover, exocytosis events are less frequent at cell borders similarly to non-patterned cells. Using the same micropatterns, we then measured membrane tension by TIRF-FLIM. Interestingly, cells displayed a radial gradient of membrane tension with lowest membrane tension values at the extreme cell border and in the center, and a linear increase in membrane tension for the center to the periphery (F4D-E). The Moran’s index is lower in patterned cells than in non-patterned cells despite the presence of the gradient (S4C). Taken together with the PLL experiments, this result points out a correlation between clustering/compartmentalization of membrane tension and clustering of exocytosis. The cross-correlation between the normalized lifetime and the exocytosis probability demonstrates that the exocytosis probability increases with the membrane tension in a monotonous however non-linear manner (F4F). Note that the exocytosis probability is defined as the probability of exocytosis at a given place knowing that an event will occur. Therefore, it does not reflect the exocytosis rate but only its spatial distribution. The above results together indicate that exocytosis is favored at regions with high local membrane tension, and that these regions are spatially organized leading to clustering in these regions.

### 5) Strength of membrane tension gradient regulates clustering of lysosomal exocytosis

To further investigate the role of membrane tension in the spatial regulation of exocytosis, we established exocytosis and membrane tension maps of cells cultured on ring-shaped micropatterns of different sizes (F5A). The average membrane tension was found to be similar on all pattern sizes (S5A). However, the membrane tension gradient varied according to the pattern size (F5B), except for cells plated on small patterns that did not display any gradient. Consistently, the Moran’s index increased from small to large micro-patterns (S5B). Interestingly, the absence of a gradient in the smallest size micropatterns correlates with a lower level of clustering of exocytosis events, whereas the presence of a gradient in the largest cell size correlates with higher clustering (F5C). On the other hand, the exocytosis rate (normalized to cell surface) was not significantly different regardless of the pattern sizes (S5C). These results confirm a role of the membrane tension gradient in the regulation of spatial exocytosis patterns.

## Discussion

Our data demonstrate that lysosomal exocytosis is not random (*i*.*e*. CSR) but clustered in space and time with a positive coupling between spatial and temporal dimensions (Fig1). Lysosomal exocytosis occurs close to internal FAs and it is almost absent at cell borders. In agreement with this result, cells seeded on PLL, which inhibits the formation of FAs, display a decreased clustering of exocytic events (Fig2). Previous work described a targeting of lysosomes to FAs but exocytosis at these points was not demonstrated (Schiefermeier et al., 2014). Our data suggest that clustering does not rely on an intact cytoskeleton as the spatial pattern of lysosomal exocytosis was not perturbed upon treatment with drugs affecting the cytoskeleton (Fig2). The underlying mechanism supporting clustering of exocytotic events from lysosomes at FA is likely different from the exocytosis of Golgi-derived vesicles. Indeed, clustering of Golgi-derived vesicles was shown to be inhibited by both actin and microtubule depolymerization (Yuan et al., 2015). However, interfering with the actin cytoskeleton inhibits VAMP7-mediated exocytosis rate in neuronal cells (Gupton and Gertler, 2010). The involvement of the cytoskeleton in clustering of lysosomal and Golgi-derived exocytosis will then require further investigation in different cell types.

**Figure 1.**
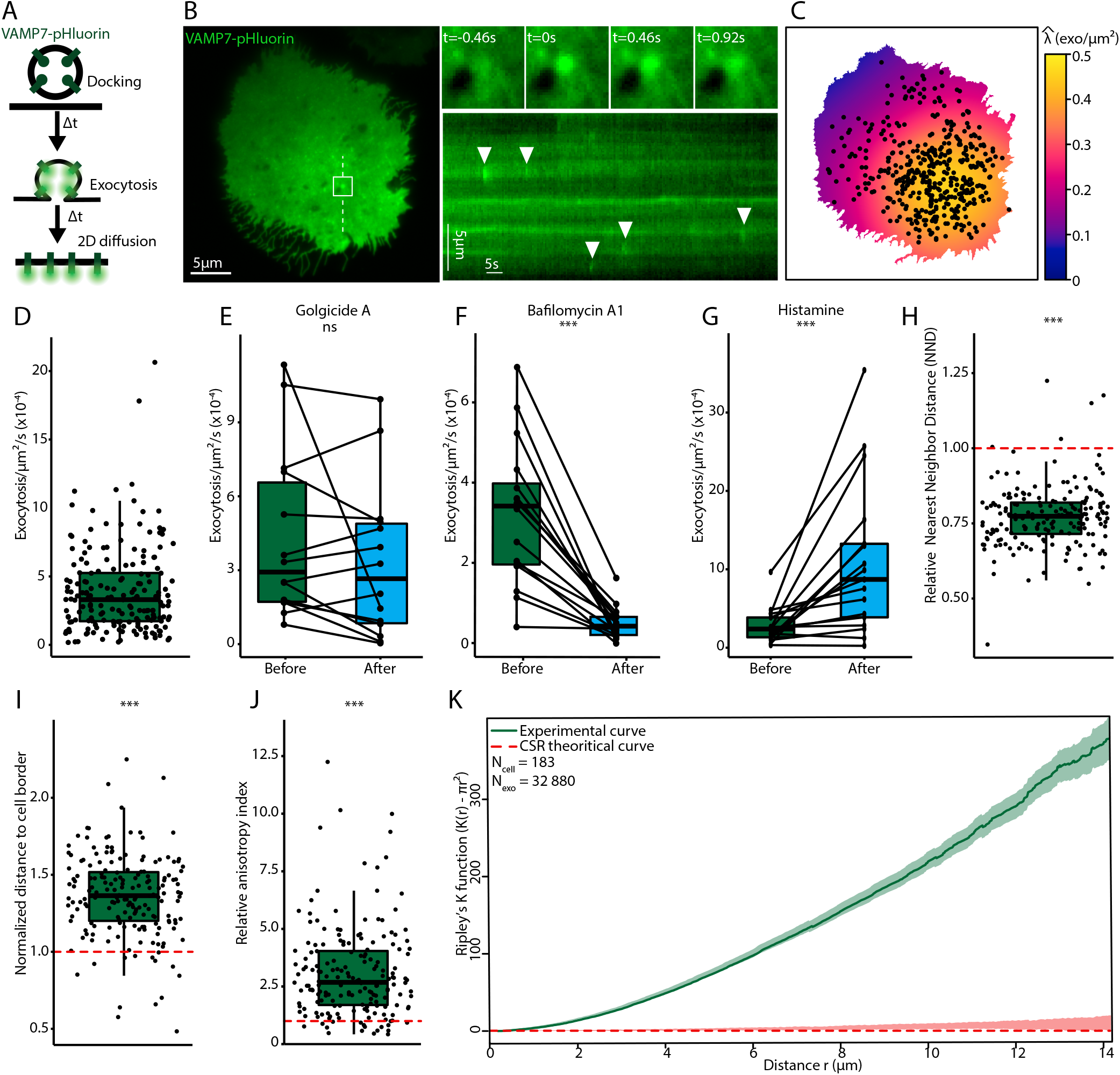
Lysosomal exocytosis is not random but clustered. **A**. Schematic representation of the exocytosis of a VAMP7-pHluorin+ vesicle: the low pH of the acidic lumen quenches the fluorescence of pHluorin. During exocytosis, protons are released and pHluorin starts to emit light. An exocytosis event is followed by the 2D diffusion of VAMP7-pHluorin at the plasma membrane. **B**. TIRFM image of VAMP7-pHluorin in a transfected RPE1 cell. The inset represents the field in the white square showing one exocytosis event at different time points, t=0 represents the beginning of the exocytosis event. A events. **C**. Exocytosis intensity map of the cell in B acquired during 5min. Black dots represent exocytosis kymograph is plotted along the dashed white line and arrowheads indicate several observed exocytosis events. The color code represents the estimation of the local intensity 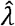 expressed in exocytosis/μm^2^. **D**. Normalized exocytosis rate in RPE1 cells from n=183 cells (and 32.880 exocytosis events) from 34 independent experiments. (3.9×10^−4^ ± 3.1×10^−4^ exocytosis/s/μm^2^). **E**. Exocytosis rate before and after Golgicide A (10μM, 30min) treatment; n=14 cells from 3 independent experiments. **F**. Exocytosis rate before and after bafilomycin A1 (100nM, 60min) treatment; n=16 cells from 3 independent experiments. **G**. Exocytosis rate before and after histamine (100μM, no incubation) treatment; n=17 cells from 3 independent experiments. **In E-G**, significance has been evaluated with paired Wilcoxon test, ns p>0.05 and ***p<0.001. **H**. Relative Nearest Neighbor Distance (NND_observed_/NND_simulated_) of basal exocytosis observed in D. **I**. Relative distance to cell borders (distance_observed_/distance_simulated_) of basal exocytosis observed in D. **J**. Relative anisotropy index (anisotropy_observed_/anistropy_simulated_) of basal exocytosis observed in D. 90% of the cells have a relative anisotropy index superior to 1. **In H-J**, the red dotted line represents expected value under CSR hypothesis and the significance of the deviation to CSR has been computed using a t-test. ***p<0.001. **K**. Average centered Ripley’s K function (K(r)-πr^2^) of 183 cells (and 32.880 exocytosis events) observed in D. Green curve represents the experimental results ± SEM and the red dotted line the expectation under CSR hypothesis. Red shade represents envelope containing 95% of CSR simulations.

**Figure 2.**
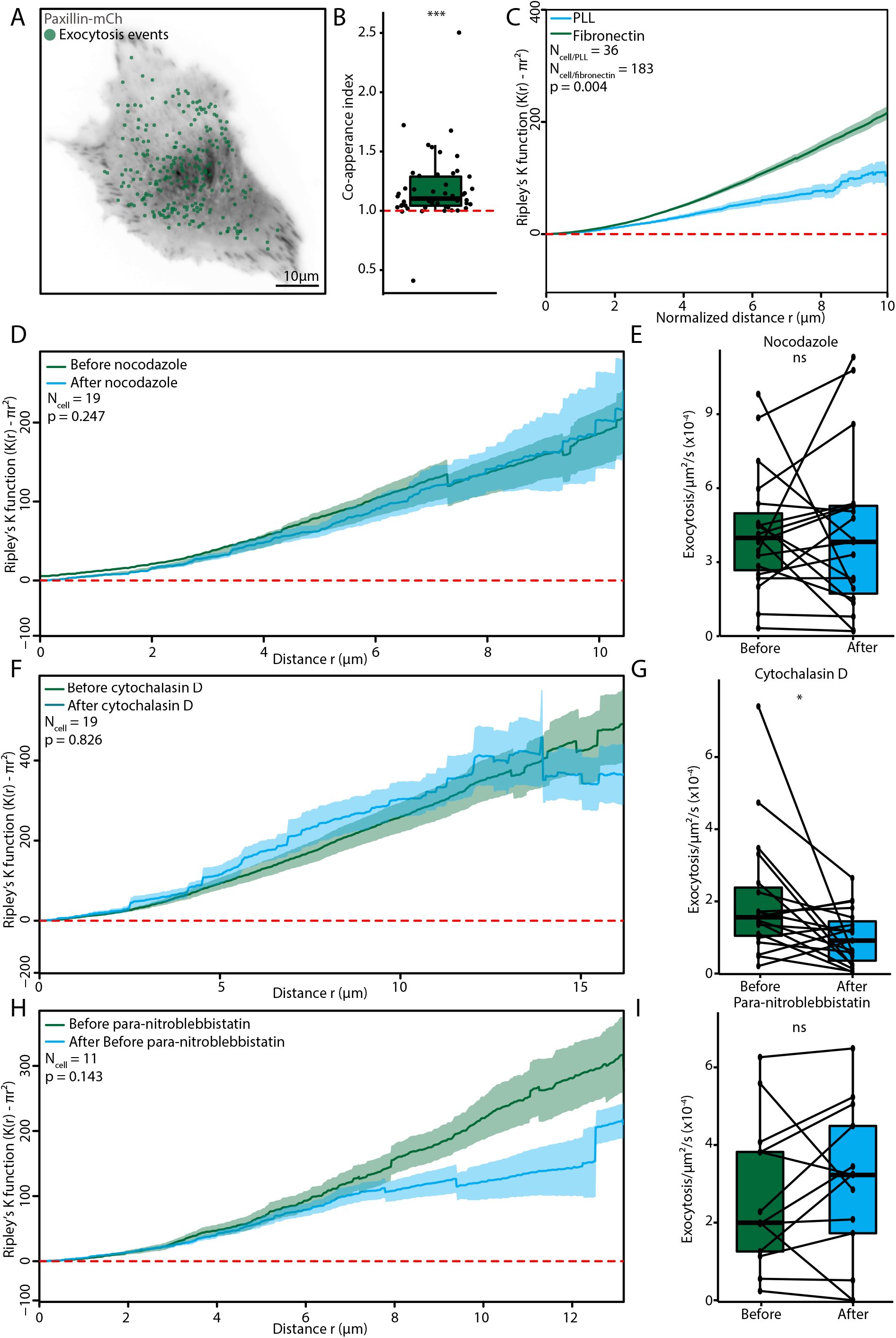
Lysosomal exocytosis is coupled to internal focal-adhesions in a cytoskeleton-independent manner. **A**. Merge from TIRFM live cell imaging of a paxillin-mCh/VAMP7-pHluorin co-transfected RPE1 cell: the gray scale image represents snapshot of paxillin-mCh intensity and green dots represent exocytosis events localization. **B**. Co-appearance index between exocytosis events and FAs. Red dashed line represents expected co-appearance in the case of CSR. The significance of the deviation to CSR has been computed using a t-test, ***p<0.001; n=49 cells from 4 independent experiments. **C**. Average spatial Ripley’s K function ± SEM for cells seeded on fibronectin (green curve) and PLL (blue curve); 36 cells on PLL and 183 cells on fibronectin were analyzed from 3 and 34 independent experiments, respectively. **D**. Average spatial Ripley’s K function ± SEM before (green) and after (blue) incubation with nocodazole (10μM, 60min). **E**. Exocytosis rate before and after nocodazole treatment (10μM, 60min). **D-E**: n=19 cells analyzed from 4 independent experiments. **F**. Average spatial Ripley’s K function ± SEM before (green) and after (blue) cytochalasin D treatment (500nM, 60min). **G**. Exocytosis rate before and after incubation with cytochalasin D (500nM, 60min). **F-G**: 19 cells analyzed from 3 independent experiments. **H**. Average spatial Ripley’s K function ± SEM before (green) and after (blue) para-nitro-blebbistatin treatment (20μM, 15min). **I**. Exocytosis rate before and after para-nitro-blebbistatin treatment (20μM, 15min). **H-I**: n=13 cells from 3 independent experiments were analyzed. In **C, D, F** and **H**, the significance of the difference between Ripley’s K functions has been evaluated using a permutation test (see methods) and red-dashed line represents expectation in the case of CSR. In **E, G** and **I**, the significance has been evaluated using a paired Wilcoxon test, ns p>0.05, *p<0.05.

We found that interfering with membrane tension by hypo-osmotic shock and methyl-β-cyclodextrin treatments impact clustering of exocytosis, revealing for the first time a role of membrane tension in the spatial organization of exocytosis (Fig3). Our results complement previous data illustrating the role of membrane tension on the exocytosis rate (Wen et al., 2016; Kliesch et al., 2017; Shi et al., 2018; Wang and Galli, 2018; Wang et al., 2018; Cohen and Shi, 2020).

**Figure 3.**
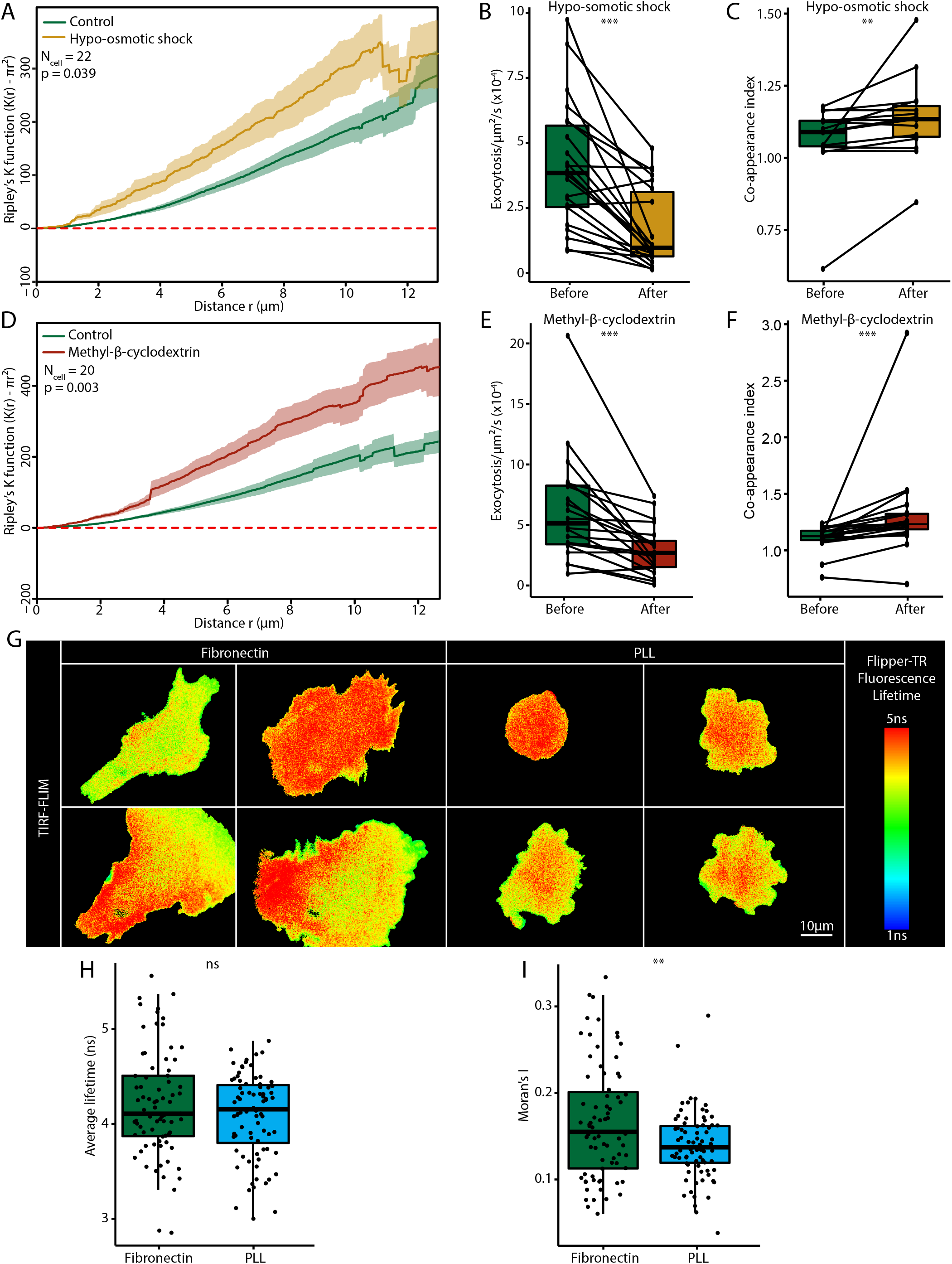
Membrane tension regulates lysosomal exocytosis clustering. **A**. Average spatial Ripley’s K function ± SEM before (green) and after (yellow) hypo-osmotic shock (1:1 dilution, 15min). **B**. Exocytosis rate before and after hypo-osmotic shock. In **A** and **B**, 22 cells from 4 independent experiments were analyzed. **C**. Co-appearance index between exocytosis spots and FAs before and after hypo-osmotic shock. 15 cells analyzed in 3 independent experiments. **D**. Average spatial Ripley’s K function ± SEM before (green) and after (dark-red) β-methyl-cyclodextrin addition (5mM, 15min). **E**. Exocytosis rate before and after β-methyl-cyclodextrin addition. In **D** and **E**, 20 cells from 3 independent experiments were analyzed. **F**. Co-appearance index between exocytosis spots and FAs before and after β-methyl-cyclodextrin; 19 cells from 3 independent experiments were analyzed. In **A** and **D**, the significance of the difference between Ripley’s K functions has been evaluated using a permutation test (see method) and red-dashed line represents expected curve in the case of CSR. In **B, C, E** and **F**, significance has been evaluated using paired Wilcoxon test, **p<0.01 and **p<0.001. **G**. TIRF-FLIM images of each 4 representative cells seeded on either fibronectin or PLL substrate, and incubated with Flipper-TR. The color code represents the Flipper-TR fluorescence lifetime. **H**. Average fluorescence lifetime per cell seeded on fibronectin and PLL substrate. **I**. Moran’s I index per cell seeded on fibronectin and PLL substrate. In **H** and **I**, significance has been evaluated using t-test, ns p>0.05 and **p<0.01, 75 cells have been analyzed on fibronectin-coated surface, and 80 cells in PLL-coated, from 3 independent experiments.

A previous study reported that in addition to changing total membrane tension, methyl-β-cyclodextrin treatment increases its heterogeneity (Biswas et al., 2019). In addition, transmembrane proteins such as FAs have been proposed to behave as obstacles to the lipid flow that equilibrates membrane tension (Cohen and Shi, 2020). Cells seeded on PLL reveal a more uniform membrane tension, confirming the role of FAs as obstacles. Together, our data suggest that an inhomogeneity of membrane tension induced by FAs leads to the accumulation of secretory events at these regions. This hypothesis is supported by experiments performed on micropatterns, which show that the symmetric arrangement of FAs leads to a well-defined gradient of membrane tension (Fig.4). Moreover, changing the micropattern diameter increases the strength of membrane tension gradient and exocytosis clustering (Fig.5). Interestingly, a gradient in membrane tension has been already observed in moving keratinocytes (Lieber et al., 2015). Such a gradient could result from the friction between plasma membrane and either actin treadmilling or the adhesion substrate (Schweitzer et al., 2014). In our experiments performed on non-migratory cells constrained by adhesion on micropatterns, only actin retrograde flow could potentially cause friction. Yet, experiments in non-patterned cells revealed that clustering is independent of the actin cytoskeleton. Thus, the above result could indicates that the diffusion of lipids in the plasma membrane is very slow, resulting in a stable gradient of membrane tension at the time scale of our experiments, similarly to what was previously observed in non-neuronal cells (Shi et al., 2018). Finally, the experiments on micropatterns revealed a positive coupling between exocytosis probability and membrane tension (Fig. 4). Of note, the absence of gradient in membrane tension observed in cells grown on small micropatterns does not totally abolish exocytosis clustering. This suggests that other mechanisms regulate clustering, such as for instance clustering of syntaxins. In conclusion, we propose that the spatial clustering of lysosomal exocytosis relies on the spatial organization of membrane tension, which is regulated by the presence and localization of FAs.

**Figure 4.**
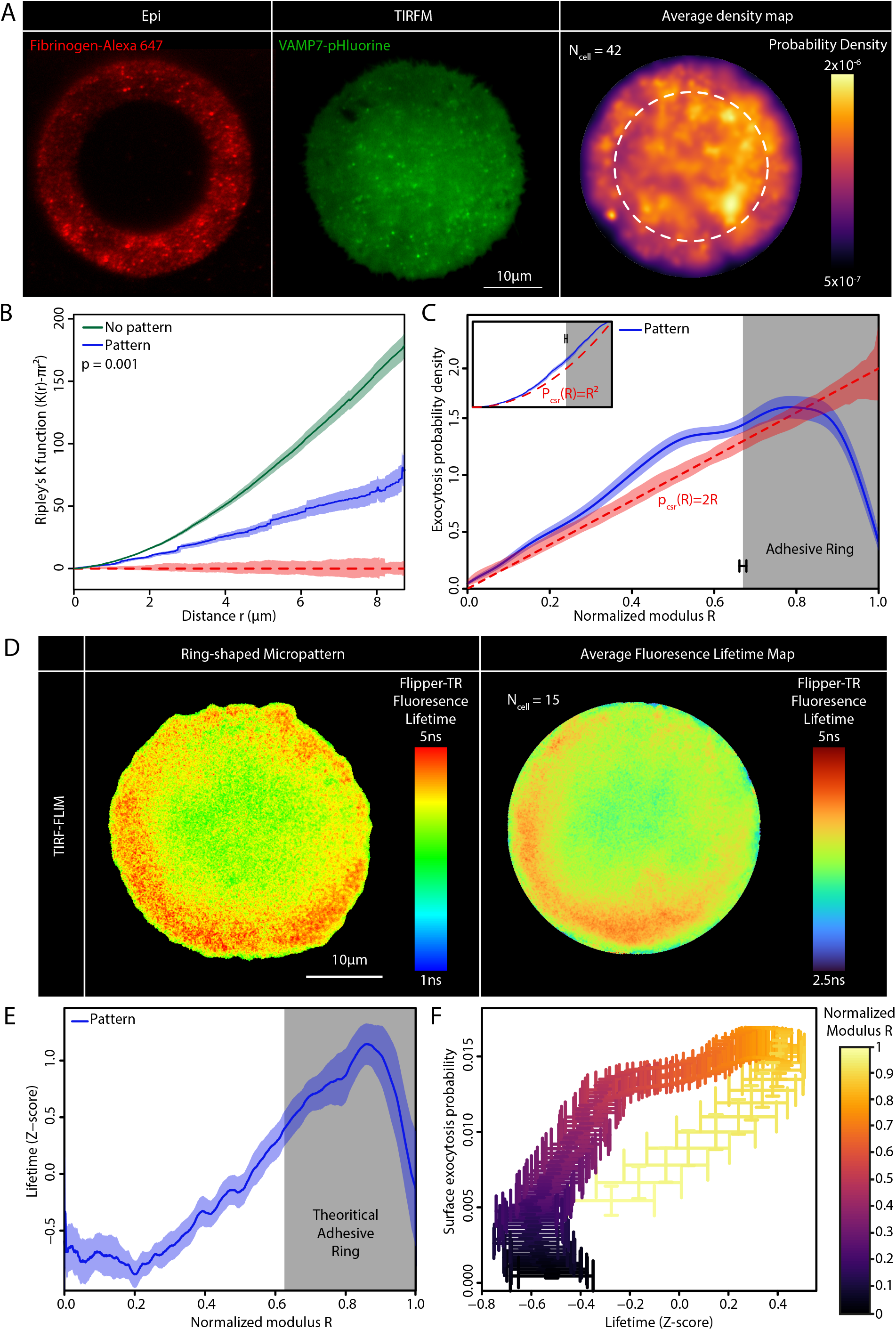
Coupling between membrane tension gradients and exocytosis probability. **A**. Representative epifluorescence image of a 37μm diameter ring-shaped micropattern stained with fibrinogene-Alexa647 and a TIRFM image of VAMP7-pHluorin transfected cell cultured on it accompanied by the average exocytosis map from 42 cells. The color code represents the probability to observe exocytosis knowing that one event occurred. **B**. Average spatial Ripley’s K function ± SEM of exocytosis in cells seeded on ring-shaped micropattern (blue) and non-patterned cells (green, same data as in F1K). The significance of the difference between Ripley’s K functions has been evaluated using a permutation test (see method). **C**. Average radial-density ± SEM of exocytosis in cells seeded on ring-shaped micropattern. The modulus (distance to pattern center) is normalized to the cell radius, setting cell border at R=1. The gray rectangle represents the average position of the adhesive part of the micropattern ± SEM. The inset represents the same data but as a cumulative density function instead of probability density. **A-C**: 42 cells were analyzed from 9 independent experiments. In **B-C**, the red-dashed line represents expected curve in the case of CSR and red shade represents envelope containing 95% of CSR simulations. **D**. TIRF-FLIM image of a representative cell seeded on a ring-shaped micropattern and incubated with Flipper-TR, and average fluorescence lifetime map from TIRF-FLIM images of 15 cells. The color code represents Flipper-TR fluorescence lifetime. **E**. Radial average ± SEM of the Flipper-TR fluorescence lifetime. Fluorescence lifetime is presented under a Z-score form (see methods) and the modulus is normalized to the cell radius setting cell border at R=1. In **D** and **E**, 15 cells analyzed in 1 independent experiment. **F**. Coupling between exocytosis probability and membrane tension using data from **C** and **E**. The estimation of exocytosis probability at a given modulus R is based on the number of events normalized by the corresponding surface, contrarily to **C**. The color code corresponds to the normalized modulus R. Each point is associated to SEM for both axes.

## Figures Legend

**Figure S1. A**. Exocytosis kinetics monitored by the normalized average fluorescence intensity (± SEM) at exocytosis event localization. Intensity has been normalized to 1 at t=0 observed at the beginning of the exocytosis event. The kinetic has been fitted by a single exponential function 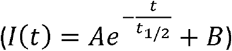 represented by the black dashed line. The half-life extracted from this fit is t_1/2_ = 1.48s. **B**. Half-life t_1/2_ measured by the same fitting but cell by cell. Average half-life is 1.69 ± 0.83s. **C**. Average cumulative exocytosis events after histamine addition: the y-axis represents the fraction of exocytosis events observed at a given time over the total number of exocytosis events recorded. Time is normalized between 0 and 1 and histamine (100μM) is added at 0.5 corresponding to the dashed black line. Green curve represents results before histamine and the blue one after histamine addition. Experimental curves are averages ± SEM; n=17 cells from 3 independent experiments. **D**. Anisotropy assay. VAMP7-pHluorin/mCh-RAB6A co-transfected cells are seeded on rectangular micropattern (visualized by fibrinogen Alexa647). Exocytosis axis was established by TIRFM and compared to the nucleus-Golgi axis. Histogram presents the percentage of cells with co-polarization and anti-polarization; n=42 cells from 4 independent experiments. Binomial test, *p<0.05. **E**. Average spatial Ripley’s K function ± SEM before (green) and after (blue) Golgicide A (10μM, 30min) addition; n=14 cells from 3 independent experiments. **F**. Average spatial Ripley’s K function ± SEM before (green) and after (blue) bafilomycin A1 (100nM, 60min) addition; n=16 cells from 3 independent experiments. **G**. Average spatial Ripley’s K function ± SEM before (green) and after (blue) histamine (100μM, no incubation) addition; n=17 cells from 3 independent experiments. **E-G**. The significance of the difference between Ripley’s K functions has been evaluated using a permutation test (see method) and red-dashed line represents expectation under CSR hypothesis. **H**. Correlation between the exocytosis rate and the clustering measured as the area under the Ripley’s K function curve between 0μm and 8μm in control cells. Correlation has been evaluated using Pearson correlation (r=0.29) and t-test (p=4.279×10^−5^) for correlation. **I**. Average Ripley’s K function by keeping 25, 50, 75 and 100% of the events for each cell (artificial thinning). Curves represent averages ± SEM and red-dashed line represents expectation under CSR hypothesis. **J**. Average centered temporal Ripley’s K function (K(t)-2t). Green curve represents the experimental average ± SEM and the red dashed line the expectation under CSR hypothesis. Red shade represents envelope containing 95% of CSR simulations. **K**. Fourier analysis of the temporal distribution of exocytosis events. Curve represents the average modulus of FFT. **L**. Median spatio-temporal Ripley’s K function (K(r,t)) subtracted by the product of the spatial and the temporal Ripley’s K function (K(r) and K(t)). An independency test has been made cell by cell. Among significant cells, 82.7% present a positive spatiotemporal coupling (and 62.8% among all cells). In **A-B** and **H-L**, 183 cells from 34 independent experiments were analyzed.

**Figure S2. A**. STX3/4 immunofluorescence of representative fixed cells seeded on ring-shaped micropattern. **B**. Average Ripley’s K function ± SEM of STX3 segmented spots from 28 cells. **C**. Average Ripley’s K function ± SEM of STX4 segmented spots from 30 cells. In **B** and **C**, red-dashed line represents expected curve in the case of CSR, and the red shade represents the envelope containing 95% of CSR simulations. **D**. Bright-field and TIRFM images of paxillin-mCh transfected cells seeded on fibronectin or PLL substrate. The cell counters of transfected cells are marked by white dashed lines. **F**. Exocytosis rate for cells seeded on fibronectin or PLL substrate. 36 cells on PLL and 183 cells on fibronectin were analyzed from 3 and 34 independent experiments, respectively. The significance has been evaluated using a t-test, ***p<0.001.

**Figure S3. A**. TIRF-FLIM images of 4 representative cells in each condition: control, hypo-osmotic shock (1:1 dilution, 15min) and β-methyl-cyclodextrin (5mM, 15min), all incubated with Flipper-TR. The color code represents the Flipper-TR fluorescence lifetime. **B**. Average fluorescence lifetime per cell in 3 conditions: control, hypo-osmotic shock and β-methyl-cyclodextrin. The significance has been evaluated using Kruskal-Wallis test and a **post-hoc** Dunn’s test, *p<0.05, 67 cells were analyzed in control condition, 43 in hypo-osmotic shock condition, and 41 cells in β-methyl-cyclodextrin, from 5, 3 and 3 independent experiments, respectively.

**Figure S4. A**. Representative epifluorescence image of a 37μm diameter ring-shaped micropattern stained with fibrinogene-Alexa488 and a TIRFM image of paxillin-mCh transfected cells. Arrowheads show internal FAs. On the right, average paxillin signal (from 19 cells) after denoising. **B**. Average paxillin radial intensity ± SEM. The modulus is normalized to the cell radius setting cell border at R=1. The intensity is normalized to obtain a total equal to 1, 19 cells were analyzed. **C**. Moran’s index for cells seeded on micropattern or non-patterned fibronectin coated classical culture, 15 cells from one independent experiment and 42 cells from 9 independent experiments were analyzed, respectively. The significance has been evaluated using unpaired Wilcoxon test, *p<0.05.

**Figure S5. A**. Average Flipper-TR fluorescence lifetimes per cell seeded on ring-shaped micropatterns with different diameters: small, medium and large. Significance has been evaluated using Kruskal-Wallis test, ns p>0.05. **B**. Correlation between the Moran’s index per cell and the cell radius (cell seeded on ring-shaped micropatterns). Correlation has been evaluated using Pearson correlation (r=0.27) and t-test (p=0.0217) for correlation. In **A** and **B**, 60 cells from 5 independent experiments were analyzed. **C**. Correlation between the exocytosis rate per cell and the cell radius (cell seeded on ring-shaped micropatterns). Correlation has been evaluated using Pearson correlation (r=-0.08) and t-test (p=0.47) for correlation. 90 cells were analyzed from 11 independent experiments.

## Materials and Methods

### Cell culture

hTERT-immortalized retinal pigment epithelial cell line (hTERT RPE-1) were cultivated in DMEM/F12 media (Gibco, catalog # 21041-025) complemented with 10% Fetal Bovine Serum (Eurobio, catalog # CVFSVF00-01) (without antibiotics). Cells were maintained at 37°C with 5% CO2 in a humidified incubator.

### Transfection

Cells were transfected with the following constructs: VAMP7-pHluorin (Chaineau et al., 2008), mCh-Rab6A and Paxillin-mCh (Fourriere et al., 2019). Cells are transfected with 800ng of DNA (or 2×400ng for co-transfection) using the JetPrime kit (Polyplus). Cells were imaged 24h after transfection.

### Drug treatments

Cells were treated with the following drugs with the given concentration and incubation time: Golgicide A 10μM 30min (Merck, catalog # G0923), Bafilomycin A1 100nM 1h (MedChemExpress, catalog # HY-100558), Histamine 100μM no incubation (Merck, catalog # H7125), Nocodazole 10μM 1h (Merck, catalog # M1404), Cytochalasin D 500nM 1h (Merck, catalog # C8273), Para-nitroblebbistatin 20μM 15min (Cayman Chemical Company, item 24171), β-methyl-cyclodextrin 5mM 15min (Merck, catalog # C4555). Hypo-osmotic shocks were made by adding water in the media with a volume ratio of 1:1 and cells were imaged 15min after. For all drug conditions, a paired design has been used: the same cell is imaged before and after the treatment.

### Micropatterning

We followed the photolithography micropatterning protocol from Azioune *et al*. (Azioune et al., 2010). Briefly, coverslips (1.5H ThorLabs, Catalog # CG15XH1) were oxidized by plasma-cleaner (Harrick Plasma) during 5min. Coverslips were PEG-coated by incubating them on a drop of PLL-g-PEG (Surface Solutions, PLL(20)-g[3.5]-PEG(2)) (0.1mg/ml diluted in water, 10mM HEPES, pH=7.4) in a moiety chamber during 1h. After coating, patterns were printed using a deep UV lamp (Jelight Company Inc, catalog # 342-220) with radiation passing through a photomask (DeltaMask) during 5min. Finally, patterns were fibronectin-coated by incubating coverslips on a drop of fibronectin (Merck/Sigma, catalog # F1141) (50μg/ml diluted in water) and fibrinogen-Alexa647 (Molecular Probes, Invitrogen, catalog # F35200) (or fibrinogen-Alexa488) (5μg/ml) in a moiety chamber during 1h. Coverslips were conserved at 4°C in PBS.

Cell seeding on micropatterns was described in Lachuer *et al*. (Lachuer et al., 2020). Briefly, coverslips were maintained in magnetic chamlides for live imaging or kept in a P6 wells for fixation. ∼200 000 trypsinized (Thermo Ficher, catalog # 12605010) cells were added in the chamlide chamber. After 10min incubation in 37°C incubator, cells were attached to the substrate. Cells were washing using DMEM/F12 media with 20mM HEPES (Gibco, catalog # 15630-056) (+ 2% penicillin/streptomycin (Gibco, catalog # 15140-122) if cells were used for lived-imaging). Cells were incubated at least 3h in the incubator until full spreading on the micropattern. Cells were imaged the same day.

Different geometries of micropatterns were used. In figure 4 and S2, ring-shaped micropatterns with a diameter of 37μm were employed. In figure 5, 3 sizes of ring-shaped micropatterns were used, with diameters of 25μm, 35μm and 45μm. For all sizes, the thickness of the adhesive ring was 7μm. Figure 5 also includes data of figure 4. Despite these theoretical sizes, a variation in the measured cell diameter was observed likely due to UV diffraction during the printing. The actual dimensions were systematically measured. During analysis, cells with diameter inferior to 20μm were categorized as “Small”, between 20μm and 38μm as “medium” and superior to 38μm as “large”. In figure S1, rectangular micropattern has dimensions of 9×40μm.

**Figure 5.**
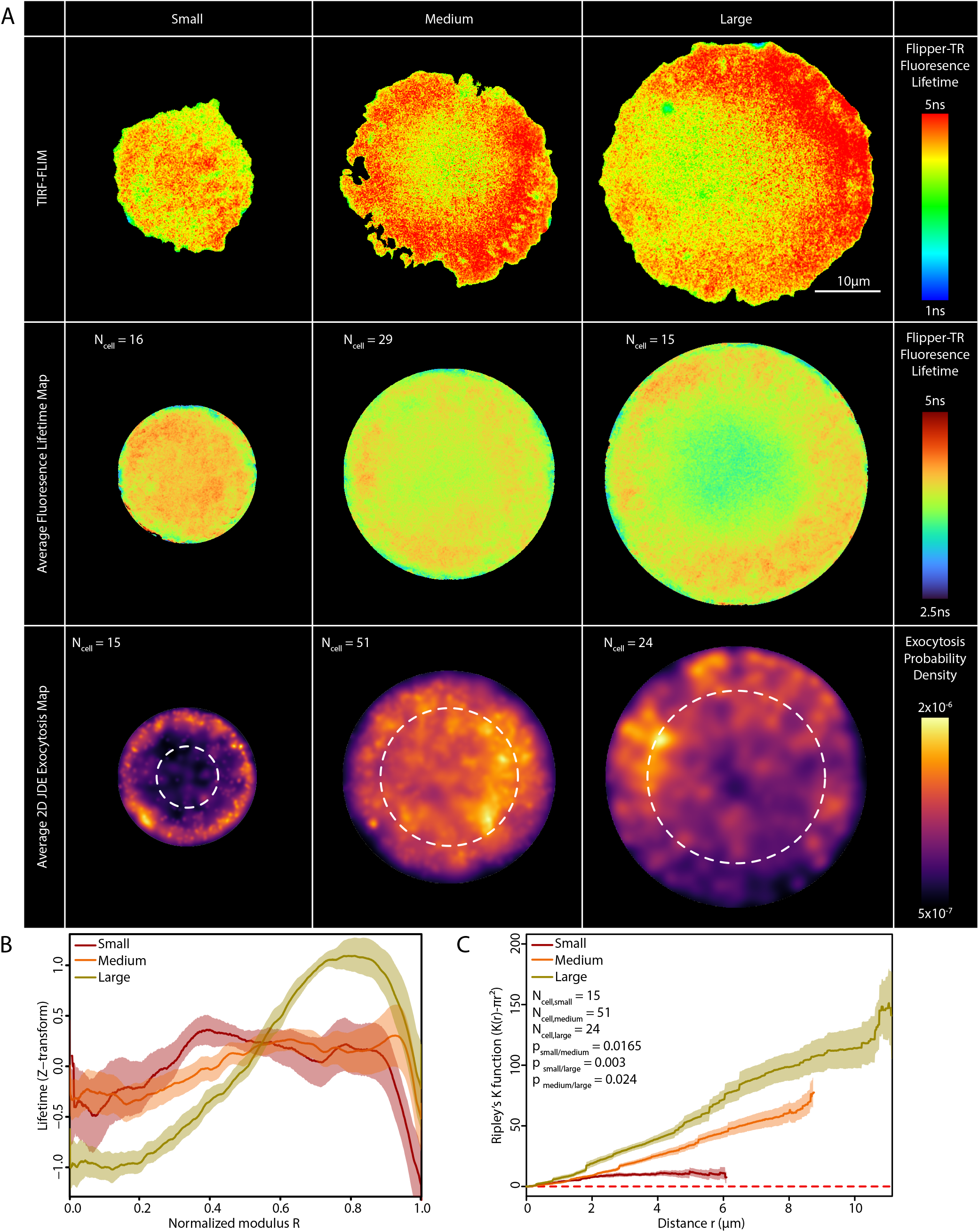
Strength of membrane tension gradient regulates clustering of lysosomal exocytosis. **A**. Representative images of cells seeded on ring-shaped micropattern of different diameters (classified as small ∼ 25μm, medium ∼ 35μm and large ∼ 45μm) and incubated with Flipper-TR visualized by TIRF-FLIM (upper panel). The color code represents the Flipper-TR fluorescence lifetime. Average fluorescence lifetime maps for cells seeded on different diameters (middle panel), maps are an average of FLIM images from n=16 (small), n=29 (medium) and n=15 (large) cells from 4 independent experiments. Color code represents Flipper-TR fluorescence lifetime. Average exocytosis maps (lower panel) from n=15 (small), n=51 (medium) and n=24 (large) cells from 11 independent experiments. The color code represents the probability of exocytosis. **B**. Radial averages ± SEM of the Flipper-TR fluorescence lifetime (data from **A**). Lifetime is presented under a Z-score form (see methods) and the modulus is normalized by the cell radius setting cell border at R=1 for the 3 conditions. **C**. Average spatial Ripley’s K functions ± SEM of exocytosis in cells seeded on ring-shaped micropatterns with different diameters, small in dark red, medium in orange and large in yellow (data from **A**). The significance of the differences between Ripley’s K functions has been evaluated using a permutation test (see methods) and p-values were corrected using Benjamini-Hochberg procedure. The red-dashed line represents expected curve under CSR hypothesis.

### Microscopy

#### TIRFM

Non-patterned cells were seeded in fluorodishes (World Precision Instrument) coated with fibronectin (or PLL (Merck P4707)). DMEM/F12 media + 20mM HEPES was used for imaging. Patterned cells were prepared as described. Acquisition was made using an inverted Nikon TIRFM equipped with an EMCCD camera (efficiency 95%) with a 100x objective (pixel size = 0.160μm). The following lasers were used 491nm, 561nm and 642nm. Time-lapse of VAMP7-pHluorin was acquired with a frame rate of 1 image every 300ms during 5min. Frame rate was set according to the half-life of exocytosis events. Due to microscopic device delay, the actual frame rate was computed using the computer time of saved files.

#### TIRF-FLIM

Images were acquired on a homemade setup based on a x100 1.49 Nikon Objective (Blandin et al., 2009; Marquer et al., 2011). A 2MHz supercontinuum laser source (SC450 HE-PP Fianium) was filtered (Excitation filter 482-18, Dichroic Di01-R488, Emission filter Long Pass 488, Semrock) within the microscope cube to match the dye excitation/emission spectra. The average power in the back focal place of the objective was between 30 and 100 μW depending on the experiment. The TIRF angle was finely controlled thanks to a motorized stage which allows one to adjust the pulsed beam focalization in the back focal plane of the TIRFM objective. Fluorescence images were detected thanks to a time-resolved detection based on the use of a high-rate imager (Kentech Ltd., UK) optically relayed to a charge-coupled device camera (ORCA AG, Hamamatsu, binning 2×2). This intensifier was synchronized with the laser pulse through a programmable delay line (Kentech, precision programmable 50 Ω delay line), which enables us to open temporal gates with 1 ns width at different times after the pulse, thus sampling the fluorescence decay. Each time gated image corresponds to an average of 10 images (10 × 250 ms). FLIM maps were thus produced by recording a series of 17 time-gated fluorescence intensity images and fitting the data for each image pixel to a single exponential decay model by use of a standard nonlinear least-squares fitting algorithm.

#### Flipper-TR

Cells were prepared as for classical TIRFM. 15min before acquisition, Flipper-TR was added in the media (Spirochrome, Catalog # SC020) (Colom et al., 2018) at a concentration of 1μM. If used, drugs were added with Flipper-TR. Cells were not washed as recommended. Due to the high variability of the fluorescence lifetime, a z-score transformation was applied when specified. For each cell, the average μ and standard-deviation σ of the fluorescence lifetime was computed. Then, each lifetime pixel x_i_ value was reduced and centered:

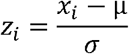

### Immunofluorescence

Cells were fixed with 4% PFA (Euromedex, catalog # 15710) during 15min and quenched with a 50mM NH4Cl solution. After PBS washing, cells were permeabilized (and blocked) with a PBS Saponin (MP Biomedicals, catalog # 102855) (0.5g/l) BSA (Merck, catalog # 10735094001) (1g/l). Coverslips were incubated during 1h in a moiety chamber at RT with primary antibodies diluted in a PBS 2% BSA solution. After PBS washing, coverslips were incubated with a secondary antibody (400x) conjugated with a fluorophore following the same protocol. Finally, coverslips were mounted with Mowiol (Biovaley, catalog # MWL4-88-25) and DAPI (Merck, catalog # D8417). The following primary antibodies were used: Syntaxin 3 (100x) (Merck/Sigma, catalog # S5547), Syntaxin 4 (1000x) (BD Transduction Laboratories, Material # 610439). The following secondary antibodies were used: Mouse A488 (Interchim 715-545-151) and Rabbit A488 (Interchim, 711-545-152).

Sample were imaged using an inverted videomicroscope with deconvolution (Delta Vision – Applied Precision) equipped with Xenon lamp. Acquisition was made at 100x (pixel size = 65nm). Images acquired were deconvolved using softworx (enhanced ratio method).

### Statistical analysis

All statistical analysis were made with R (R Core Team (2021)) with the help of the following packages: spatsat (Baddeley et al., 2015), raster, viridis, ggplot2, dunn.test, ape (Paradis and Schliep, 2019), imager, pracma, circular, ggpur, evmix (Hu and Scarrott, 2018), splancs, OpenImageR, minpack.lm.

#### Hypothesis testing

The number of cells and the number of independent repetition is indicated in the legend. Since our cells are mostly isolated when imaged, we performed only single cells analysis, each cell is considered as independent, setting the sample size. The statistical test used is indicated in the legend. Tests are always conducted in a two-sided manner and a multiple comparison correction is applied when needed. We mainly used non-parametric test (Wilcoxon test, Kruskal-Wallis test), and a parametric test (Student t-test) only when the sample size was high (n>30). Paired tests were used for all experiments where the same cell is imaged before and after treatment. Finally, correlation was measured by Pearson correlation coefficient and tested with a t-test.

#### Intensity map

The intensity function λ is defined by:

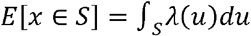

With × a pattern of points, u coordinates, S a region of the observation window, and E[x∈n] the expected number of points in S. The intensity map was computed using spatstat function density() with Jones-Diggle improved edge correction. The intensity map can be interpreted and normalized as a density map by dividing it by the total number of points.

#### Monte-Carlo CSR simulations

Monte-Carlo simulations were used for 3 purposes: i) normalization and ii) generation of CSR 95% envelopes.

1. Several measures used depend on the cell geometry and the number of exocytosis events. Therefore, we normalized observed values by CSR simulated values. For each cell, a high number (n=100) of Monte-Carlo simulations was run to generate CSR exocytosis maps associated with the same number of exocytosis events and the same cell geometry. The ratio between observed and average simulated values allows classifying cells in two categories: more extreme or less extreme than CSR compared to the observed measure.
2. Monte-Carlo simulations were used to generate CSR envelopes. Some measures (mainly Ripley’s K function) were averaged over a population of cells; therefore CSR simulations were run in the same way: Monte-Carlo CSR simulations were run for the full population of cells and the evaluated quantity is averaged over the different simulated cells. This procedure was repeated a high number of times (n=100). The CSR envelope contains 95% of these simulations.

#### Nearest Neighbor Distance

The Nearest Neighbor Distance (NND) of a given exocytosis event is the distance to the closest exocytosis event. These distances give information on the short scale spatial structure. NND was used to test CSR hypothesis according to the procedure presented in Lachuer *et al*. (Lachuer et al., 2020) and detailed in the Monte-Carlo section. Simulated average NND are compared to the observed NND. This allowed to classify the cell as clustered (NND_observed_<NND_simulated_), or dispersed (NND_observed_>NND_simulated_). Note that temporal NND were treated similarly to spatial NND, just by reducing the dimension of the analysis.

#### Ripley’s K function

For a point pattern X={x_1_, x_2_, …, x_n_} where each × is point coordinates observed in an area |**S**|, the Ripley’s K function (Ripley, 1976; Dixon, 2014) is defined as:

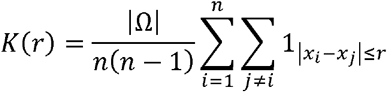

This function quantifies the average number of points in a disk of radius r centered on one point. In case of CSR, this function should be close to πr^2^. Therefore we always substrate πr^2^ to K(r), a positive value indicates clustering whereas a negative value indicates dispersing. The spatial Ripley’s K function was computed using spatstat function Kest() with the best edge correction possible. Ripley’s K function was computed between 0 and a quarter of the cell size to avoid edge effects. All Ripley’s K functions plotted are an average of a population of Ripley’s K functions (+/-SEM). A permutation test (with 999 permutations) based on a Studentized distance is used to compare populations thanks to the sptatstat function studpermu.test() (Hahn, 2012). Finally, a CSR 95% envelope was computed with Monte-Carlo simulations (see corresponding section).

Temporal Ripley’s K function K(t) is the reduction of the 2D spatial Ripley’s K function in 1D. Therefore the expected value under CSR hypothesis is 2t. K(t) was computed by our own function using the Ripley’s edge correction following (Yunta et al., 2014). Temporal Ripley’s K function was treated in the same way as the spatial one to generate the CSR 95% envelope and for the averaging.

Spatio-temporal Ripley’s K function K(r,t) is the 3D extension of the spatial Ripley’s K function that evaluates the number of neighbors in a cylinder of a radius r and a half height t (Diggle et al., 1995). K(r,t) was computed using the splancs function stkhat(). The median K(r,t) was computed (averaging is avoided due to the generation of aberrant values by splancs). In absence of spatio-temporal coupling (*i*.*e*. independency of the temporal and spatial point coordinates):

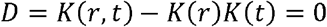

Thereby the independency can be evaluated by the D statistics. It can be statistically tested using a permutation test (1000 permutations) using the splancs function stmctest().

#### Distance to cell border

Distance to cell border was computed using imager function distance_transform(). For each cell, the distance to cell border at exocytosis sites was measured. The distribution was computed using Kernel Density Estimation (KDE). The distribution depicted is a distribution averaged over multiples cells +/-SEM. The 95% CSR envelope was computed by Monte-Carlo simulations (see corresponding section). The solid blue line represents the average of the simulated densities.

The significance at the single cell level was accessed using Monte-Carlo simulations (see corresponding section). Observed cell border distances were divided by the average simulated cell border distances allowing a classification of the cells into two categories: borders-avoiding (ratio<1) or borders-linking (ratio>1).

#### Anisotropy

The cell was cut in 30 angular sections (from the center of mass). An angle θ_i_ was associated to each section. The number of exocytosis event was computed in each section and divided by the surface of the corresponding section giving a coefficient w_i_ (normalized by the sum of the coefficients). The anisotropy/polarization measurement is based on the average resultant length computed as:

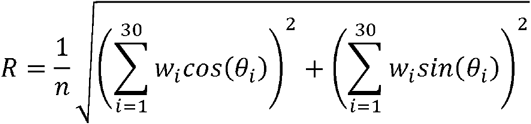

This polarization index ranged between 0 and 1. The significance was accessed using Monte-Carlo simulations (see corresponding section). Observed average resultant lengths were divided by the average simulated resultant length allowing a classification of the cells into two categories: polarized (ratio>1) or non-polarized (ratio<1).

#### Exocytosis-FA co-appearing index

The average paxillin fluorescence intensity was measured at the locations of exocytosis event. The significance was accessed using Monte-Carlo simulations (see corresponding section). The comparison of the observed average intensity with simulated ones allows classifying cells with a colocalization index (I_obs_/I_sim_) under two categories: co-appearing (I_obs_/I_sim_>1), no-co-appearing (I_obs_/I_sim_<1).

#### Modulus distribution

On micropatterns, exocytosis events were described with polar coordinates (with the origin at the center of the pattern). The modulus is the distance between an event and the center (normalized by the cell radius). For each cell a modulus distribution was computed using Kernel Density Estimation (KDE). To avoid boundary effects, an asymmetric beta kernel was used (Chen, 1999) using the evmix function dbckden(). The depicted modulus distribution is an average over the population +/-SEM. This distribution can be compared to the expected distribution in case of CSR:

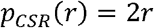

The 95% CSR envelope was computed using Monte-Carlo simulations (see corresponding section). Despite beta edge correction, the simulations do not fit perfectly p_csr_. In order to avoid any bias due to kernels, we also conducted the same analysis using empirical cumulative distributions. The empirical cumulative distribution P_CSR_ under CSR hypothesis is:

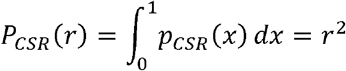

#### Fourier analysis

Fourier analysis was conducted using Fast Fourier Transform function fft(). The spectrum of modulus was computed for each cell between 0 and Nyquist frequency. Modulus spectrum were averaged over the cell population.

#### Moran’I

Moran’s I evaluates the spatial auto-correlation (Moran, 1950). We used it to evaluate the spatial structure of lifetime measurement. For an image of N pixels, the Moran’s I is defined as:

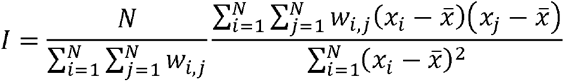

With x the i^th^ pixel value, 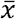 the average pixel values, and w the inverse of the Euclidian distance between pixel i and j. This index is superior to -1/(N-1) in case of pixel clustering and inferior to in case of regular spacing (*i*.*e*. pixels with similar values are regularly separated). A decreased of the Moran’s I should be perceived as a spatial decorrelation *i*.*e*. pixels are less clustered. Moran’s I index was computed using ape function Moran.I() using a random subsample of pixels (Monte-Carlo random sampling scheme) to keep a decent computation time.

### Image analysis

#### Exocytosis detection

Exocytosis events were detected manually (because automatization failed to reach our exigency level). However some events were probably missed (false negative). This is not a problem, because Ripley’s K function is invariant under random thinning (Baddeley et al., 2015). It is also likely that some events did not correspond to exocytosis (false positive). Due to the superposition principle (Baddeley et al., 2015), the experimental Ripley’s K function is the sum of the exocytosis Ripley’s K function and the one of false annotation. Therefore, under the assumption that these false annotations are random, possible false annotations could only slightly underestimates clustering.

#### Half-life

Fluorescence intensity was measured at exocytosis event localizations in a window of 1.12μm centered on the event through time. The intensity was divided by the intensity at the beginning of the event normalizing the exocytosis peak at 1. The intensity profile of all events in a cell was averaged. The intensity was fitted on this averaged profile over 20 frames (∼8s) with a single exponential function:

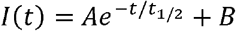

With t_1/2_ the half-life. The fitting was performed with the minpack.lm function nlsLM(). The profile depicted is the average over multiple cells. The half-life obtained from this average curve is close to the lifetime averaged over multiple cells.

#### Segmentation

Cell segmentation to obtain mask or segmentation of syntaxin patches was perforled using manual thresholding with ImageJ software.

## Acknowledgements

We are grateful to Sabine Bardin, Pallavi Mathur, David Pereira, Tarn Duong, Mathieu Coppey and Pierre Sens for fruitful discussions and/or help at the bench. We also thank Mathieu Piel lab for sharing photomasks, Thierry Galli for the VAMP7-pHluorin plasmid and syntaxin antibodies, and Marc Tramier and Giulia Bertolin for testing Flipper-TR with their FAST-FLIM. Finally, we thank (again) Thierry Galli and Christophe Lamaze for critical reading of the manuscript. The authors greatly acknowledge the Nikon Imaging Centre @ Institut Curie-CNRS, member of the French National Research Infrastructure France-BioImaging (ANR10-INSB-04). This work was supported by ARC (Association pour la Recherche sur le Cancer) PhD fellowship and FRM (Fondation Recherche Médicale) PhD extension fellowship.

